# An EEG study of Detection without Localisation in Change Blindness

**DOI:** 10.1101/513697

**Authors:** Catriona L. Scrivener, Asad Malik, Jade Marsh, Michael Lindner, Etienne B. Roesch

## Abstract

Previous studies of change blindness have suggested a distinction between detection and localisation of changes in a visual scene. Using a simple paradigm with an array of coloured squares, the present study aimed to further investigate differences in event-related potentials (ERPs) between trials in which participants could detect the presence of a colour change but not identify the location of the change (*sense* trials), versus those where participants could both detect and localise the change (*localise* trials). Individual differences in performance were controlled for by adjusting the difficulty of the task in real time. Behaviourally, reaction times for *sense*, *blind*, and *false alarm* trials were distinguishable when comparing across levels of participant certainty. In the EEG data, we found no significant differences in the visual awareness negativity ERP, contrary to previous findings. In the N2pc range, both awareness conditions (*localise* and *sense*) were significantly different to trials with no change detection (*blind* trials), suggesting that this ERP is not dependent on explicit awareness. Within the late parietal positivity range, all conditions were significantly different. These results suggest that changes can be ‘sensed’ without knowledge of the location of the changing object, and that participant certainty scores can provide valuable information about the perception of changes in change blindness.

## Introduction

Change Blindness is a phenomenon in which changes to a visual scene are often missed (Rensink, 2004; Simons & Levin, 1997). To manipulate this in an experimental setting, the change blindness paradigm typically consists of two images displayed in quick succession that are interrupted by a blank screen or a distractor image. In some instances, the second image is identical to the first, and in others, some aspect will have changed. Participants are then asked to report if the trial contained a change or not. The complexity of these images varies across paradigms, ranging from coloured rectangles (Koivisto & Revonsuo, 2003) and coloured dots (Schankin & Wascher, 2007), to facial expressions (Eimer & Mazza, 2005), detailed visual scenes (Fernandez-Duque et al., 2003) and household objects (Busch et al., 2010). In all cases, although complete visual information is available, participants often fail to notice or identify changes.

Most versions of the change blindness paradigm ask participants to detect the presence of a change across two image presentations, meaning that trials can only be categorised as one of four types: hit (or *see* trials), miss (or *blind* trials), false alarm (FA), or correct rejection (CR), depending on whether the participant reports seeing a change. Several researchers have challenged the traditional view that vision must always be accompanied by a complete conscious visual experience, or the activation of complete internal representation of what we see (Rensink, 2004; Fernandez-Duque & Thornton, 2000), and subsequently suggested the possibility of further trial divisions in the change blindness paradigm. In an early experiment reported by Rensink (2004), participants were asked to indicate when they ‘thought’ that something had changed in a flicker paradigm, and again when they were certain that they could see the change. In a flicker paradigm, the original image and changed image are presented sequentially until the participant is able to detect the change. Trials in which these responses had a time difference greater than 1 second were labeled as trials with a ‘significant duration of sensing’, where the participant suspected a difference but was not confident in their perception of the change. Rensink (2004) termed the ability to detect a change without fully identifying it as *sensing*, suggesting that this condition is both phenomenologically and perceptually distinct to the traditionally reported *see* condition.

Several other researchers have explored the possibility of an awareness condition that lies somewhere between the traditional *see* and *blind* dichotomy (Fernandez-Duque et al., 2003; Laloyaux et al., 2006; Thornton & Fernandez-Duque, 2001; Galpin et al., 2008; Busch et al., 2009; Ball & Busch, 2015; Kimura et al., 2008; Hollingworth et al., 2001). For example, Fernandez-Duque et al. (2003) found that the location of a change could be identified above chance level even when participants did not report to see the change itself (but see Mitroff et al. 2002 and Laloyaux et al. 2006 for a discussion of these results). Further, in Mitroff et al. (2004) participants were able to identify pre- and post-change object stimuli above chance level when they detected a change, as well as when they did not. The presence of a *sense* condition has therefore been suggested as evidence that change blindness may arise from a failure to compare two displays or images, rather than a failure to encode the visual information (Simons & Rensink, 2005; Hollingworth et al., 2001). Further, *sense* trials may occur when features of a changing object only reach a pre-attentive stage, and are not fully integrated at later stages of visual processing (Galpin et al., 2008; Busch et al., 2009).

Results from change blindness experiments using EEG appear to support this assertion. In previous EEG research, the trials types of *see* and *blind* are often distinguishable in an early visual attention component around 200-300 ms after the change onset at contralateral electrode sites, known as the N2pc (Luck & Hillyard, 1994; Schankin & Wascher, 2007), as well as a late central parietal component at 400-600ms, known as the late parietal positivity (Railo et al., 2011; Koivisto & Revonsuo, 2003). There is also evidence that the amplitude of early visual components, such as P1 and N1, may be dependent on the awareness level of the participant during a change detection task (Pourtois et al., 2006; Railo et al., 2011). The presence of an N2pc reflects the allocation of attention towards an attended object (Luck & Ford, 1998), and the amplitude is increased for ‘aware’ stimuli (Schankin & Wascher, 2007). While the N2pc has also been found for ‘unaware’ stimuli in a masking paradigm, and therefore does not necessarily represent conscious awareness of a change (Woodman & Luck, 2003), another ERP called the visual awareness negativity (VAN) is thought to be dependent on spatial attention and conscious report (Koivisto et al., 2008). Similarly, the LPP is thought to reflect conscious aspects of task processing (Railo et al., 2011), and has been shown to correlate with participants’ confidence in their responses (Eimer & Mazza, 2005).

Several EEG papers have also identified differences between *see*, *sense* and *blind* conditions. In a comparison between trials in which the participants were able to detect a change and identify the object of the change (*see*), and those where they could detect a change but not name it (*sense*), Busch et al. (2010) found an increase in amplitude of the visual awareness negativity, or VAN, which is thought to be dependent on spatial attention (Koivisto et al., 2008). The same effect was found in a later P3/LPP ERP at posterior electrodes. However, the N2pc peak was found only when participants could both detect and identify the change, and was not present when participants were change blind, or could not identify the object. The authors concluded that *seeing* a change is not simply a stronger version of *sensing* a change, as the N2pc can be found for *see* trials and but not *sense* trials. This supports the hypothesis of Rensink (2004) that *seeing* and *sensing* may be facilitated by separate mechanisms. Other studies have also found differences in ERP amplitudes when comparing trials with ‘full’ and ‘partial’ awareness (Fernandez-Duque et al., 2003; Kimura et al., 2008; Busch, 2013; Ball & Busch, 2015), but the definition of *sense* trials and ‘full’ awareness varies across studies (Mitroff et al., 2002), leading to divergent results.

The main aim of the present study was to compare behavioural and ERP effects for trials in which participants could report the presence of a change but not localise it (*sense*), versus those in which participants could report and localise the change correctly (*localise*). Specifically, we divided the visual display into quadrants, and asked participants to select the quadrant in which the change occurred. Our *sense* condition therefore requires registration of the change, but not necessarily knowledge of its location (Mitroff et al., 2002). Further, participants were asked to rate how confident they were in their responses at every trial, in order to distinguish between trial types (Galpin et al., 2008). We used a simple paradigm with an array of coloured squares (see figure 1).

**Figure 1.**
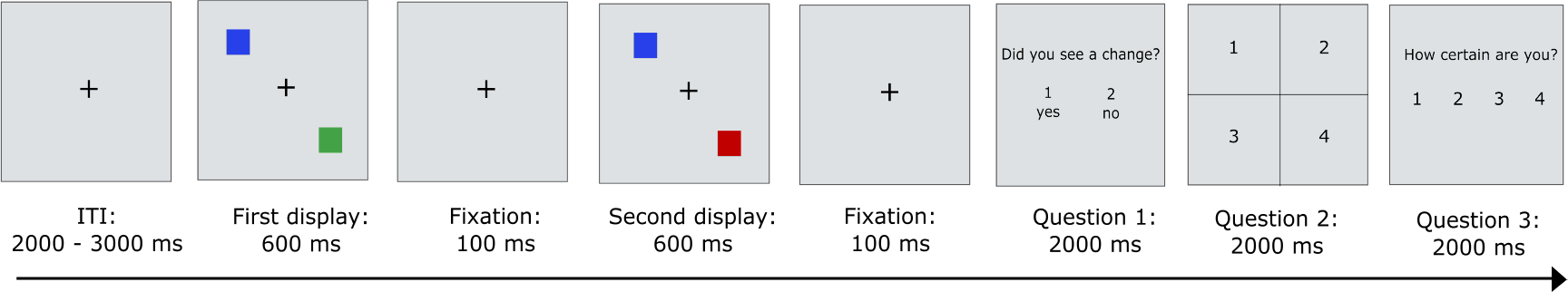
Illustration of the experimental paradigm. The number of squares presented varied from 2 to a maximum of 36. Question 1 asked ‘Did you see a change?’ to which participants could respond ‘Yes’ or ‘No’. Question 2 asked participants to localise the change, based on a grid from top left to bottom right. Question 3 asked how certain participants were of their responses, ranging from ‘1: Very Uncertain’ to ‘4: Very Certain’. If participants responded ‘no change’ to question 1, they were moved straight on to question 3.

As increased amplitudes in the N2pc and LPP have previously been found in the *see* condition compared to the *blind* condition, we hypothesised that we would replicate these findings (Railo et al., 2011). Although modulation of P1 amplitudes have been reported in some change detection paradigms (Busch et al., 2009; Pourtois et al., 2006), others report no such effect (Eimer, 2000; Turatto et al., 2002; Niedeggen et al., 2001), so our hypothesis was not directed. When comparing *localise* versus *sense* trials, we hypothesised that we would find increased amplitudes in the VAN, LPP, and N2pc for *localise* trials (Busch et al., 2010; Fernandez-Duque et al., 2003).

A further aim of the study was to identify if *sense* trials are behaviourally different to *blind* or *false alarm* trials, as others have suggested (Fernandez-Duque et al., 2003; Galpin et al., 2008), or whether they can be explained by explicit mechanisms (Mitroff et al., 2002). If the *sense* condition can be explained by participant pressing the incorrect response when they did not see a change, then reaction times for *sense* trials should be similar to *blind* trials. Or, if *sense* can be explained by a liberal response criteria, such that participants report seeing a change despite not being sure, then uncertain *sense* trials should have similar reaction times to *false alarms*. By using EEG measures of neural activity, as well as additionally asking participants to rate their confidence at each trial (Galpin et al., 2008), we aimed to distinguish between these distinct types of awareness.

## Materials and methods

### Participants

Twenty subjects (mean ± SD, age = 20 ± 5, 6 left handed, 2 male) with no history of psychiatric or neurological disorders participated in this EEG study. All had corrected-to-normal vision and were not colour blind (based on self report). The experiment was approved by the University of Reading ethics committee (UREC: 17/03), and was conducted in accordance with the Declaration of Helsinki (as of 2008). All participants gave informed consent to take part, including consent to share their anonymised data. Three participants were removed from the original sample size of 23 for having less than 200 usable trials after pre-processing (out of a maximum of 250 trials). Trials were classified as unusable if they contained muscle or eye-movement artifacts that could not be removed during pre-processing.

### Stimuli and Presentation

Participants were presented with a change blindness task using Psychtoolbox (Kleiner et al., 2007), on a 1920 × 1080 monitor with a 60 Hz refresh rate. Participants were seated comfortably on an armchair, at approximately 60cm away from the screen, alone, in a quiet room (Faraday cage) with constant dim light. They were asked to fixate on a central fixation cross and identify changes between consecutive displays of coloured squares. These were interrupted by a short fixation display to facilitate the change blindness phenomenon (see figure 1 for details on display durations). On change trials, one of the squares changed colour from the first to the second display. On no-change trials, the displays were identical. This was followed by two or three questions, depending on the participant’s response to the first question. Each participant completed 5 blocks of 50 trials, leaving a total of 250 trials. Within these trials, two-thirds contained a change in coloured square (165 trials), and the rest contained no change (85 trials).

Question 1 asked ‘Did you see a change?’ to which participants could respond ‘yes’ or ‘no’ using a keyboard. Question 2 asked participants to localise the change, based on a 2×2 grid from top left to bottom right. Question 3 asked how certain participants were of their responses, ranging from ‘1: Very Uncertain’ to ‘4: Very Certain’. If participants responded ‘no’ change to question 1, they were moved straight to question 3. This decision was made as our hypotheses did not relate to ‘implicit’ change detection, as reported in Fernandez-Duque & Thornton (2000), and removing this question allowed for a greater number of trials within the same period of time. Participants were asked to respond within a limit of two seconds for each question, and trials with any response missing were not included in further analysis (3.6 ± 2.9). Participants made their response on a keyboard, using their index finger and middle finger on each hands.

Difficulty was modulated in real time by adding and removing two squares from the display, based on the assumption that more distractors increases task difficulty (Vogel et al., 2005). This was to prevent floor and ceiling performance during the task as a result of individual differences (Luck & Vogel, 2013), and optimise for performance rather than to establish specific individual thresholds. Performance over the previous two trials was used to update the current trial; two consecutive correct answers added two squares, two incorrect deducted two squares, and one correct and one incorrect resulted in no change. The decision to increase or decrease the number of squares was made using responses to the localisation question (Q 2), as we were specifically interested in controlling the number of *sense* and *localise* trials. The number of squares always changed by two, to balance the number on the left right hemifields of the screen. The location of the change on each trial was random, but the change occurred an equal number of times on the left and right hemifield of the screen. As the colour of the squares was not related to our main hypotheses, we used seven default MATLAB colours; blue, cyan, yellow, green, white, red, and magenta (MathWorks, Inc., version 2016b).

### Behavioural Analysis

The trials in which a change occurred were divided into three conditions: *blind* (no change detection), *localise* (change detection and localisation), and *sense* (change detection without localisation). Trials in which no change occurred were divided into *correct rejection* (no change reported) and *false alarm* (change incorrectly reported). The number of false alarm trials was low, with a mean of 12.45 trials (range = 2 − 33*, SD* = .65), and therefore EEG analysis comparing *false alarm* to *sense* trials was not possible. The percentage of *false alarm* trials was calculated in relation to the the total number of no-change trials, whereas the percentage of *sense* trials was calculated in relation to the total number of change trials.

Detection accuracy for each participant was calculated based on the percentage of change trials in which they correctly detected a change. Localisation accuracy was calculated as the percentage of correctly detected changes where the localisation was also correct. We also recorded each participant’s mean and maximum difficulty scores, with the maximum referring the the highest number of squares that were displayed to them during the experiment.

D’prime was calculated as a measure of participant response bias. This was calculated using the equation *d* = *z*(hit rate) − *z*(false alarm rate) (Stanislaw & Todorov, 1999), and is defined as the difference between the means of signal and noise distributions, normalised by the variance. Response bias, or criterion, was also calculated, where *c* = −0.5 * (*z*(hit rate) + *z*(false alarm rate)) (Stanislaw & Todorov, 1999). *c* = 0 indicates no response bias to either ‘yes’ or ‘no’ responses. *c* > 0 indicates a bias towards ‘no’ responses, with fewer hits and fewer false alarms. *c* < 0 indicates bias towards ‘yes’, with more hits but also more false alarms. We expected that participants would display a range of response strategies.

One problem faced in identifying a *sense* condition is that it is difficult to distinguish these trials from *false alarm* trials, or those where participants press the wrong response key (Simons & Rensink, 2005; Mitroff et al., 2002). Rensink et al. (2004) found that reaction times for *sense* trials were shorter for change trials than no-change trials, meaning that participants were slower when they were simply making a false alarm. Galpin et al. (2008) also found greater certainty associated with *sensing* during change trials, compared to *false alarms*. We therefore compared reaction times across awareness conditions, as well as between levels of confidence. As trial numbers were low, ‘very uncertain’ and ‘uncertain’ responses were combined, and ‘certain’ and ‘very certain’ were combined. Each awareness condition therefore had two levels of certainty; for example, *localise certain* and *localise uncertain*.

### EEG Data Acquisition

EEG data was recorded with a BrainVision EasyCap (Brain Products), with 64 passive electrodes including an IO channel, arranged according to the 10-20 layout. The reference electrode was placed at FCz and the ground at AFz. Impedance was kept below 10k for all the EEG channels, and 5k for the IO channel. EEG signals were recorded using BrainVision Recorder (Brain Products, version 1.20) at a sampling rate of 5000 Hz.

### EEG Pre-processing

Raw EEG data was pre-processed using BrainVision Analyzer (Brain Products, version 2.1). The data was first downsampled to 500 Hz to reduce computation time, then filtered with a high-pass filter of 0.01 Hz to remove low frequency drift (Butterworth, 2nd order). A low-pass filter of 50 Hz and a notch filter of 50 Hz were chosen to remove line noise. Independent component analysis (ICA) was used to remove eye movement artifacts (FastICA). Two components were removed for each participant; one corresponding to eye-blinks and the other to lateralised eye-movements.

Further analysis was completed using EEGLab (Delorme & Makeig, 2004). Trials were marked as outliers if any ERP value was greater than 3 standard deviations from the mean value of that ERP across all trials (using the MATLAB function ‘isoutlier’). Note that we only searched for outliers in the electrodes used for analysis (P07, P08, Cz, Pz, and CPz). Trials marked as containing outliers were excluded from further analysis (3.25 trials per participant ± 2.46), as well as those where a response to any question was not made within the response time (3.60 trials per participant ± 2.94).

Segments were then taken from −200 to 7000 ms to include the whole trial, and baseline corrected using a mean of the data within −200ms to 0ms, where 0ms was the start of the first display of coloured squares (see figure 1). We chose the baseline period to be before the first display onset, rather than the second, as we were interested in visual ERPs that occurred in response to the both displays. It has also been suggested that ERPs in response to the first presentation of a stimuli are related to the subsequent perception of change (Pourtois et al., 2006).

### EEG Analysis

To identify the peaks of the visually evoked potentials (P1 and N1), a grand average ERP was calculated across all conditions and participants, as advised in Luck & Gaspelin (2017), from electrodes P07 and P08. From here, the peaks of interest were determined by identifying the local maxima/minima of the expected peaks, using the peak detection function in BrainVision Analyzer. The mean value within a window around the peak was used instead of the peak value, as the mean is more robust against noise (Luck, 2014). A window of 40ms around the mean was chosen as the appropriate window for visual ERPs P1 and N1. In relation to the first display onset, the first P1 was identified at 122ms, and the first N1 at 212ms. In relation to the second display onset, the second P1 was identified at 114ms, and the second N1 at 222ms.

Based on previous literature (Busch et al., 2010; Tseng et al., 2012; Fernandez-Duque et al., 2003), the N2pc was defined as the mean within 200-400 ms after the second display at occipital electrodes PO7 and PO8. Over central parietal electrodes Cz, CPz and Pz, the VAN was defined within a window of 130-330 ms after the second display, and the LPP within a window of 400-600ms. We used window sizes of 200 ms, defined a-priori, in an attempt to be conservative given the large variation within the literature.

To assess how differences between early visual components across detection conditions were reflected at each stimulus presentation, P1 and N1 amplitudes were compared in two separate 2×3 repeated measures ANOVAs, with display (first/second) and awareness (*blind/localise/sense*) as the independent variables. Differences across hemispheres in the N2pc were analysed with another 2×3 repeated measures ANOVA, with the independent variables of hemisphere (contralateral/ipsilateral) and awareness (*blind/localise/sense*). Amplitudes of the VAN and the LPP were compared in two separate repeated measures ANOVAs with awareness (*blind/localise/sense*) as the independent variable. Where Mauchly’s Test of Sphericity indicated that the assumption had been violated, Greenhouse-Geisser correction was used. All post-hoc comparisons were two tailed, and corrected for multiple comparisons using false discovery rate where *q* = .05 (Benjamini & Hochberg, 1995). Effect sizes are reported as partial eta squared for ANOVA, and repeated measures Hedge’s g for t-tests (Lakens, 2013).

To determine if the visual ERPs (P1 and N1) varied as a function of the task difficulty (the number of squares presented per trial) we correlated the single-trial P1 and N1 amplitudes with the number of squares presented at each trial. To determine if the LPP amplitude varied with participant confidence, as previously suggested (Eimer & Mazza, 2005), single-trial LPP values were correlated with participant confidence ratings. For single-trial analysis, time courses were constructed for each participant from the single-trial values of each ERP, at each channel (7 ERPs, 64 channels, 20 participants). Each single-trial value was calculated as the mean amplitude within the pre-defined ERP window at each trial. These values were baseline corrected by subtracting the mean of the trial from which they were selected. P-values were corrected for multiple comparisons using false discovery rate where *q* = .05 (Benjamini & Hochberg, 1995).

## Behavioural Results

### Accuracy and Difficulty

Accuracy for question 1, in which participants had to identify a change, had a mean of 49% (range = 32 − 73%, SD = 13). Accuracy for question 2, in which participants had to localise the change, had a mean of 70% (55 − 87%, 8). The mean difficulty level given to each participant was 14 squares (10 − 18, 3), with the mean maximum difficulty experienced by each participant at 26 squares (20 − 36, 4). D’prime scores had a mean of .61 (.74 − 1.64,.27). In a one-sample t-test, D’prime was significantly different from zero, suggesting that participants were able to distinguish between change and no-change trials *t*(19) = 19.293, *p* < .001. Two participants had a negative criterion, meaning that they had a response bias towards false alarms. All other participants had positive criterion, indicating a conservative response strategy (.60 ± .42).

Mean difficulty did not correlate with detection accuracy (*r* = −.022, *p* = .928), location accuracy (*r* = .136, *p* = .566), or d’prime (*r* = −.229, *p* = .332), suggesting that the difficulty of the task did not influence task performance. Maximum difficulty also did 6 not correlate with detection accuracy (*r* = .067, *p* = .779), location accuracy (*r* = −.077, *p* = .748), or d’prime (*r* = −.148, *p* = 535).

### Comparison of *sense* and *false alarm* trials

The percentage of *false alarm* trials (14.64% ± 11.35) was lower than the percentage of *sense* trials (30.31% ± 8.02) *t*(19) = −7.107, *p* < .001, *g_rm_* = 1.48, suggesting that *sense* trials occurred more often than participants made false alarms. However, the percentage of false alarms was positively correlated with the percentage of *sense* trials (*r* = .527, *p* = .017). Therefore, participants with a more liberal response strategy who made more false alarms, also had more *sense* trials.

Reaction times for *sense* and *false alarm* trials were compared, to determine if *sense* trials were different to trials where the participant incorrectly reported a change during a no change trial. Reaction times for all *sense* trials (0.744 ± 0.149 s), regardless of certainty, were not significantly different to *false alarm* trials (0.778 ± 0.179 s), *t*(19) = −1.229, *p* = .234, *g_rm_* = 0.193. However, *sense certain* trials (0.619 ± 0.133 s) were significantly faster than *false alarm* trials, *t*(19) = −4.741, *p* < .001, *g_rm_* = 0.939. Therefore, when participants were certain that a change occurred, they responded more quickly than when they were simply making a false alarm.

Reaction times for *sense certain* trials (0.619 ± 0.133 s) were also significantly faster than *false alarm uncertain* trials (0.817 ± 0.211 s), *t*(19) = −4.510, *p* < .001, *g_rm_* = 1.081. However, this may be explained by the general finding that, across all conditions, certain trials (.628*s* ± .142) were faster than uncertain trials (0.849 ± 0.129 s), (*t*(19) = −7.831, *p* < .001, *g_rm_* = 1.563)

### Comparison of *sense* and *blind* trials

Reaction times for *sense* trials (0.744 ± 0.149 s) were not significantly different to *blind* trials (0.731 ± 0.176 s), *t*(19) = −.285, *p* = 779, *g_rm_* = .082. However, reaction times for *sense uncertain* trials (0.801 ± 0.189 s) were significantly slower than *blind* trials, (*t*(19) = 4.424, *p* < .001, *g_rm_* = .373). Therefore, on trials where the participant did not see the change (*blind*), they responded more quickly than when they suspected a change but could not provide additional information about it (*sense*).

Comparatively, reaction times for *sense certain* trials (0.619 ± 0.133 s) were significantly faster than *blind uncertain* trials (0.860 ± 0.231 s), (*t*(19) = 4.424, *p* < .001, *g_rm_* = 1.224), which again may be explained by the fact that uncertain trials were slower over all conditions.

### Comparison of *blind* trials and no-change trials

Out of the 20 participants included in the analysis, 15 were slower to respond when they were *blind* to the change, compared to no-change trials (75%). This difference in reaction times was not significant when comparing all no-change trials (0.704 ± 0.167 s) to *blind* trials (0.731 ± 0.176 s), (*t*(19) = −2.084, *p* = .051, *g_rm_* = .143). However, *blind uncertain* trials (0.860 ± 0.231 s) were significantly slower than no-change trials (0.704 ± 0.167 s), (*t*(19) = 3.637, *p* = .002, *g_rm_* = .718). Therefore, despite being *blind* to the change, the presence of a change in the display increased reaction times, particularly for trials where the participant was uncertain.

## EEG Results

### Single-trial Correlations

The purpose of this analysis was to check whether single-trial ERPs varied as a function of difficulty, i.e. the number of squares presented on the screen during each trial. After correcting for multiple comparisons using FDR correction (*q* = .05), only one test was significant. This was a negative correlation between N2pc amplitude and difficulty at electrode POz for participant 18, which had the highest correlation result across all tests, *r* = −.3351. POz, however, was not used in the further N2pc analyses as electrodes P07 and P08 were chosen a-priori.

The second analysis was to test whether single-trial ERPs varied with the confidence ratings of the participants. Several researchers have suggested that ERPs, particularly those in later time windows such as the LPP, may be more influenced by participant confidence in their response than by the level of conscious awareness (Koivisto et al., 2005; Eimer, 2005). None of the tests were significant, with all *p* > .34. This result suggests that confidence ratings were not directly correlated with single-trial ERP amplitudes.

### P1 and N1

Overall, no significant differences were found between the three awareness conditions for either the P1 or N1 (figure 2). For P1 amplitudes, the main effect of awareness was not significant, *F* (1.473, 19) = 1.117, *p* = .338, *η*^2^ = .056. The main effect of display was also not significant, *F* (1, 19) = .355, *p* = .558, *η*^2^ = .018, nor was the interaction between awareness and display, *F* (1.80, 34.35) = .307, *p* = .305, *η*^2^ = .060.

**Figure 2.**
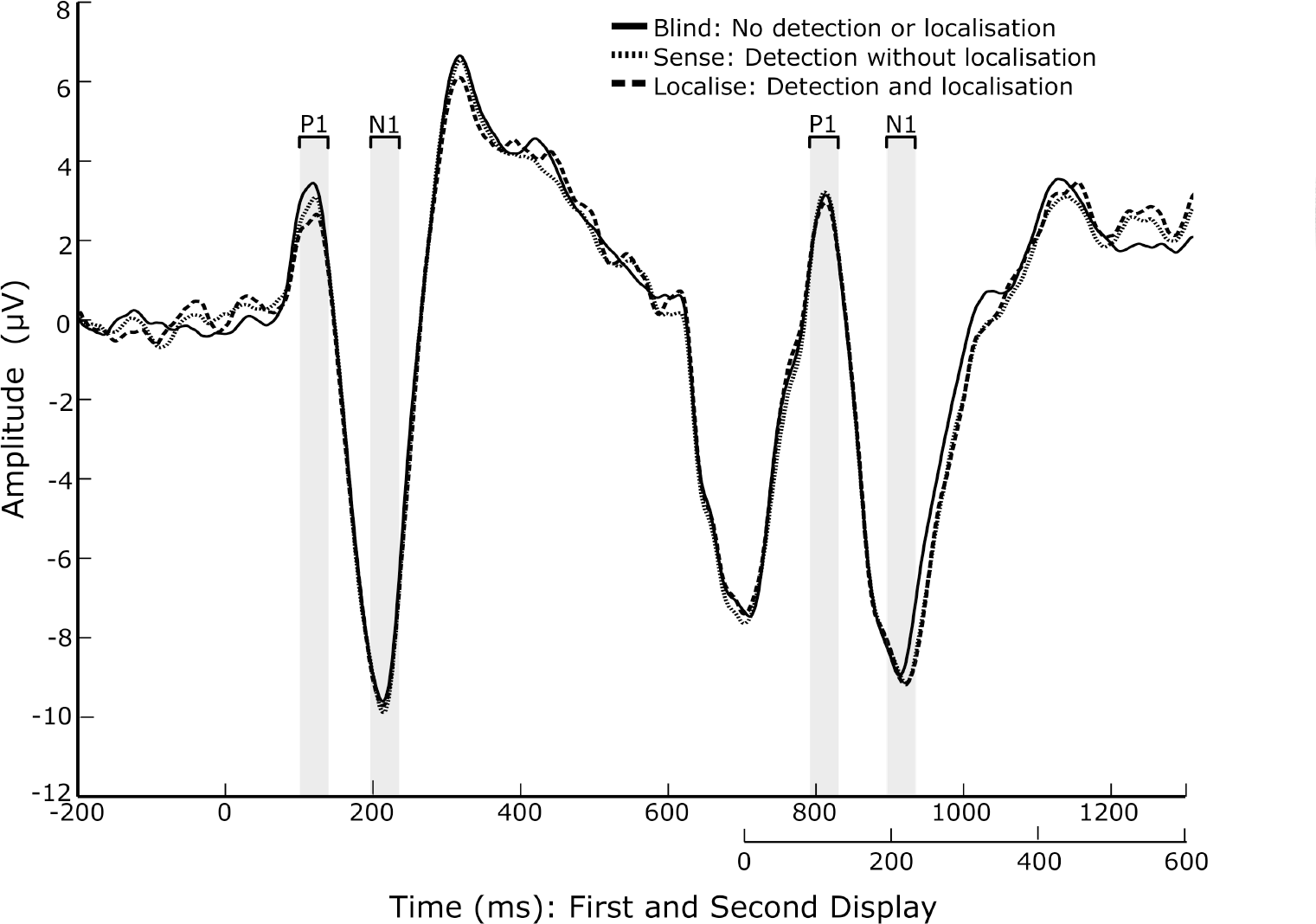
ERP plot showing the mean of electrodes PO7 and PO8, for each awareness condition. Condition means for the values within the shaded time windows were used for ERP analysis.

For the N1, the main effect of awareness was not significant, *F* (1.36, 19) = 3.534, *p* = .060, *η*^2^ = .157. The main effect of display was also not significant, *F* (1, 19) = .209, *p* = .653, *η*^2^ = .011, nor was the interaction between awareness and display, *F* (1.87, 35.61) = .377, *p* = .675, *η*^2^ = .019.

### N2pc

In line with our hypothesis, there was a significant main effect of awareness on N2pc amplitudes, *F* (2, 18) = 4.043, *p* = .026, *η*^2^ = .175 (figure 3). There was also a significant main effect of hemisphere, *F* (1, 19) = 4.594, *p* = .045, *η*^2^ = .195, with a greater negativity in the contralateral hemisphere (−2.89 ±3.97 *μV*) than the ipsilateral (−2.33 ±4.26 *μV*). The interaction was not significant, *F* (2, 18) = 1.048, *p* = .361, *η*^2^ = .052.

Post-hoc pairwise comparisons across awareness levels with a FDR corrected threshold of *p* = 0.03 showed that *blind* (−2.055 ±1.23 *μV*) had a significantly smaller N2pc amplitude than localise *localise*, (−2.941 ±1.80 *μV*), *t*(19) = 2.340, *p* = .030, *g_rm_* = .197, and *sense* (−2.847 ±1.19 *μV*), *t*(19) = 2.525, *p* = .021, *g_rm_* = .181. However, *sense* and *localise* were not significantly different, *t*(19) = −.283, *p* = .780, *g_rm_* = .022.

**Figure 3.**
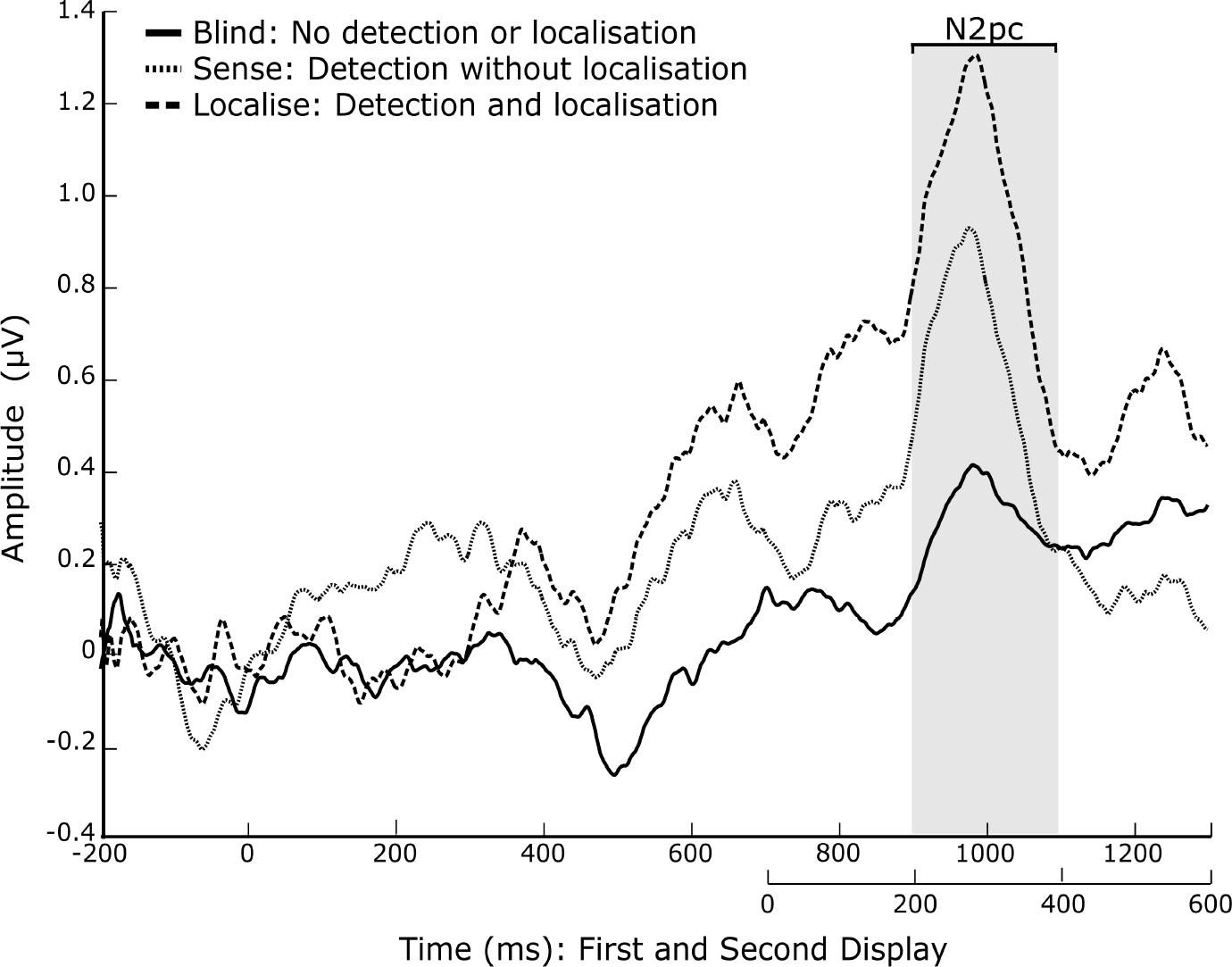
ERP plot showing a mean of electrodes PO7 and PO8, for each awareness condition. Asymmetry was calculated by subtracting contralateral from ipsilateral waveforms. Condition means for the values within the shaded time window (200-400 ms after the second display) were used for N2pc analysis.

### Visual Awareness Negativity (VAN)

Confirming our hypothesis, there was a significant main effect of awareness on the VAN (figure 4), *F* (1.374, 18) = 3.931, *p* = .046, *η*^2^ = .171. However, in post-hoc pairwise comparisons across awareness levels with a FDR corrected threshold of p = 0.04, *blind* (−1.474 ±2.52 *μV*) was not significantly different to *localise* (−2.167 ±3.09 *μV*), *t*(19) = 2.158, *p* = .044, *g_rm_* = .217, or *sense* (−1.961 ±1.92 *μV*), *t*(19) = 1.950, *p* = .066, *g_rm_* = .161. *Localise* and *sense* were also not significantly different, *t*(19) = 1.235, *p* = .232, *g_rm_* = .062.

**Figure 4.**
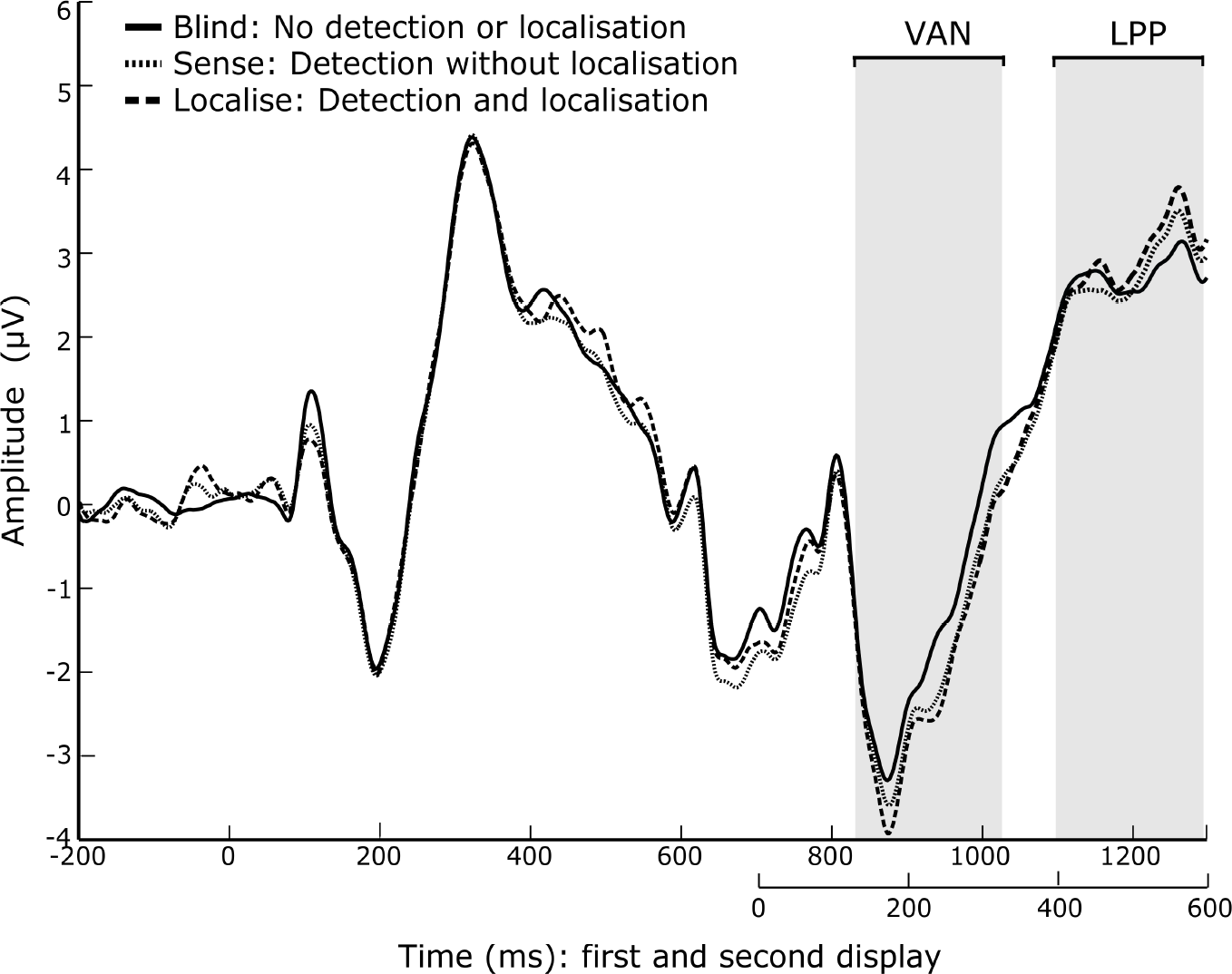
ERP plot showing a mean of electrodes Cz, CPz, and Pz, for each awareness condition. Condition means for the values within the shaded time window were used for ERP analysis. The first shaded area was used for the visual awareness negativity (130-330 ms after the second stimulus), and the second shaded area was used for the late parietal positivity (400-600 ms).

### Late Parietal Potential (LPP)

In support of our hypothesis, there was a significant main effect of awareness on LPP amplitudes (figure 4), *F* (1.355, 8) = 7.000, *p* = .008, *η* 2 = .269. In post-hoc pairwise comparisons across awareness levels with a FDR corrected threshold of p = .048, *blind* (2.931 ±2.02 *μV*) was significantly smaller in amplitude to both *localise* (3.905 ±2.53 *μV*), *t*(19) = −3.094, *p* = .006, *g_rm_* = .383, and *sense* (3.591 ±2.40 *μV*), *t*(19) = −2.193, *p* = .041, *g_rm_* = .275. *Localise* was also significantly greater in amplitude than *sense*, *t*(19) = 2.110, *p* = .048, *g_rm_* = .118.

## Discussion

The main aim of this change blindness experiment was to distinguish between trials in which participants could both detect and localise a change in coloured square (*localise*), versus those in which they could only detect it (*sense*), or not detect it at all (*blind*). We found significant differences between *blind* trials and both *sense* and *localise* trials in the N2pc ERP. We also found that *sense* and *localise* were significantly different in the late LPP window. Behaviourally, reaction time results allowed us to distinguish *sense* trials from *false alarm* and *blind* trials, when taking participant certainty into account. Overall, our results suggest that the *sense* condition may be distinguishable from the traditional *see* condition, and that utilising participant confidence is a valuable method to distinguish between levels of awareness in change blindness.

### EEG

Our results indicated a difference between *sense* and *localise* trials within the LPP range, which were significantly different to each other, as well as to *blind*. An increased late positivity for change detected trials versus change *blind* trials is the most commonly reported finding within the EEG literature, and all of the papers considered in the review by Koivisto et al. (2010) report this finding. This may be due to the relatively large size of this ERP, peaking anywhere between 300 and 700ms after a change stimulus and across large time windows.

While the earlier negativity, VAN, is typically thought to be associated with phenomenal consciousness, the later positivity is linked to reflective consciousness and greater subject report ability. The repeated finding that the LPP can be significantly reduced by specific stimuli, such as non-targets and repeated stimuli, suggests that it is not a direct correlate of visual awareness (Koivisto & Revonsuo, 2010). Instead, it is generally thought to reflect higher level or fully conscious aspects of task processing (Railo et al., 2011; Koivisto & Revonsuo, 2003). It has also be shown that the LPP correlates with confidence in participant responses (Eimer & Mazza, 2005). However, when correlating single trial LPP amplitudes with confidence ratings, we did not find a significant effect.

The majority of change blindness papers listed by Koivisto (2010) reported enhanced negativity in the N1-N2 range (with the exception of (Fernandez-Duque & Thornton, 2000; Niedeggen et al., 2001). Busch et al. (2010) found that an N2pc was evoked only when the change was fully identified, and not in the *sense* or *blind* conditions. Based on this, they draw the conclusion that for *sense* trials, the change did not induce a shift in attention towards the location of the change, and therefore the features of the change were not available for further recognition. This is based on the assumption that the N2pc represents the allocation of attention towards the object of interest, which is supported by a number of previous studies (Luck & Ford, 1998).

Contrary to this, we found that both awareness conditions were significantly different to *blind* trials, indicating a shift in the allocation of attention for all identified changes, regardless of subsequent success/failure to localise. It may also be that *sense* trials elicited a shift in attention to the correct hemifield of change (and therefore subsequently an N2pc), but that it was not specific enough to determine whether the change occurred in the upper or lower field within that hemifield. Woodman & Luck (2003) also identified an N2pc for ‘unaware’ stimuli which were masked by object substitution masking, suggesting that the N2pc does not necessarily represent conscious awareness of changes (Woodman & Luck, 2003). It is suggested, however, that the amplitude is increased for ‘aware’ stimuli (Schankin & Wascher, 2007), which our findings support.

Other studies have reported a larger N2pc for more attention-demanding tasks (Luck & Hillyard, 1994). It was therefore a concern before analysis that *sense* trials would occur more often when the task was more difficult, and therefore that the N2pc would be larger for this condition as a result of uneven trial distribution. We found the opposite, however, with a smaller N2pc in the *sense* condition compared to the *localise* condition. We also found no significant correlation with the number of *sense* trials and the difficulty of task given to the participant, suggesting that the trial distribution was even enough to avoid this confound.

Although there was a main effect of awareness within the VAN at central parietal sites, the corrected post-hoc tests were not significant, and only *localise* was significantly different to *blind* using an uncorrected threshold (*p* = .044). In comparison, (Busch et al., 2010) were able to identify a VAN for their *sense* condition, compared to *blind*. The VAN is thought to be dependent on spatial attention, and requires both the location and identity of an object to be stored such that it is available for conscious report (Koivisto et al., 2008). As participants were not able to identify the location of change in our *sense* condition, this may explain the lack of significant VAN ERP. In another study (Koivisto et al., 2008), VAN was found to be reduced when participants were asked to keep their eyes fixated at the centre of the screen. This was the case in this experiment, which may also have contributed to the lack of significant finding withing the VAN window.

Unlike previous findings from Pourtois et al., the amplitude of the P1 during the first stimuli display was not influenced by the level of awareness (Pourtois et al., 2006). In fact, no significant modulations of awareness were identified within either of the visual ERPs, P1 and N1, across either display, which fails to support previous findings that P1 amplitude during a visual display varies with attention (Wilenius & Revonsuo, 2007) and identification of changes (Mathewson et al., 2009). One possible reason for this could be that the number of squares varied across trials, unlike other experiments where the number was fixed (Pourtois et al., 2006), and therefore possibly driven by inter-individual differences in performance. However, when correlating single trial P1 and N1 amplitudes with difficulty across time, no significant correlations were found, after correcting for multiple comparisons. This suggests that the amount of squares presented during each trial had no direct influence on the amplitude of the P1 and N1, and therefore that it did not create an obvious confound in the data.

In a review of the ERP correlates of visual awareness, Koivisto & Revonsuo (2010) list a number of change blindness EEG studies that also failed to detect modulation of an early P1 peak (Eimer, 2000; Koivisto & Revonsuo, 2003; Fernandez-Duque et al., 2003; Schankin & Wascher, 2007; Turatto et al., 2002; Niedeggen et al., 2001), compared to two studies which did (Busch et al., 2010; Pourtois et al., 2006). One criticism of the change blindness paradigm is that success relies on the participant paying attention to the first visual display, in order for the change to be integrated into the short term memory and the change detected (Simons & Levin, 1997). Attention levels, and perhaps ERPs, in response to the first display, may therefore have a large influence on the success of the following trial. We did not find any electrophysiological evidence for this occurring, as the amplitude of the P1 and N1 during the first visual display did not correlate with subsequent ERPs, or with performance. It may be, however, that this effect presented itself in a section of the EEG that was not analysed, or that the effect was not strong enough to detect across participants, some of whom may have been more vigilant than others.

### Behavioural

One explanation for the presence of a *sense* condition in change blindness is that it reflects a liberal response criteria, such that participants report seeing a change even though they were not certain that it occurred (Simons & Rensink, 2005). In other words, they make a *‘false alarm’* during change trials. If this is the case, then these trials may be similar in number to *false alarm* trials, where participants incorrectly report a change for identical displays where they could not have seen a change. We found that participants had a significantly higher percentage of *sense* trials than *false alarm* trials, suggesting that *sense* trials occurred more often. This finding cannot be explained by the fact that more trials contained a change, as the percentages were calculated in relation to the total number of change/no-change trials, respectively.

However, we also found a significant correlation between the percentage of *sense* and *false alarm* trials, suggesting that participants with a more liberal response strategy were more likely to report the presence of a change when they were not completely sure where the change occurred. To further compare *sense* and *false alarm* trials, we also examined reaction times. Although all *sense* trials combined were not significantly different to *false alarms*, *sense certain* trials were significantly faster. Therefore, *sense* trials where the participant was certain that they saw something change may be distinguishable from simple *false alarms*.

Another explanation for the *sense* condition is that it contains trials for which the participant mistakenly reported a change, even though they were not aware of it. In this case, reaction times for *sense* trials should be similar to those for *blind* trials, particularly those where participants were uncertain of their responses. We found that *sense uncertain* trials were significantly slower than *blind* trials, suggesting that participants took longer to respond to trials where they suspected that something had changed, but were uncertain.

Previous studies have also reported that participants responded ‘no change’ more quickly for no-change trials, compared to change trials (Williams Simons, 2000; Mitroff, 2002). The participant’s response is the same in both trial types, but the presence of a change is different. This suggests that even when they fail to detect the change in a change trial, they take longer to respond. We therefore compared reaction times for no-change trials and *blind* trials. Out of the 20 participants, 15 were slower to respond when they were blind to the change, compared to no-change trials (75%), which is higher than the 68% reported by Williams & Simons (2000). Although no significant differences were found between all *blind* and no-change trials, *blind uncertain* trials were significantly slower. It is possible that in *blind certain* trials, no information about the change is registered by the participant, and therefore reaction times are similar to no-change trials. However, in *blind uncertain* trials, some information may be available to the participant, leading to slower reaction times, but not enough for them to be confident to report the change.

As the average accuracy for question 1 (yes/no) was roughly 50% across participants, change trials were fairly equally divided into *see* (all trials where a change was correctly identified) and *blind* conditions. Within the *see* trials, accuracy for question 2 (‘where did the change occur?’) was roughly 70%, leaving more trials in the *localise* condition than the *sense* condition. Unfortunately, the number of false alarm trials was low, meaning that a comparison of false alarms trials in the EEG data was not possible. Within the *sense* trials, there was also a low number of ‘certain’ trials, meaning that dividing the awareness conditions into certain/uncertain for EEG analysis was also not possible. Future experiments could focus on obtaining higher trial numbers, which would hopefully facilitate this analysis. However, the very nature of the *sense* condition means that participants are unlikely to be ‘certain’ during many of the trials.

We defined the difficulty of the task as the number of squares that were presented to the participant during each trial. Participants ranged in the difficulty within which they could perform the task with similar accuracy. The maximum difficulty ranged from 10 to 36, with only one participant reaching the highest possible level. The fact that the difficulty measures, such as maximum difficulty and mean difficulty, were not correlated with accuracy or d’prime, suggests that the difficulty modulation managed to control for individual differences in ability across participants. However, despite the difficulty modulation, the range of accuracy demonstrated by the participants was large (32% - 73%). Future studies could benefit from a more sophisticated measure of trial-by-trial adaptation, to further balance the number trials within each condition and participant.

## Conclusions

Overall, the main aim of this experiment was to identify neural differences between full and partial awareness of colour changes, while controlling for individual differences in performance. Behaviourally, reaction time results allowed us to distinguish *sense* trials from *false alarm* and *blind* trials, when taking participant certainty into account. For EEG data in the N2pc range, *localise* and *sense* were both significantly different to *blind* trials, but not significantly different from each other. In comparison, within the LPP range, all conditions were significantly different, indicating that the difference between levels of awareness was represented in this late potential. Overall, our results suggest that the *sense* condition may be distinguishable from the traditional *see* condition, and that utilising participant confidence is a valuable method to distinguish between levels of awareness in change blindness.

## Acknowledgements

Thank you to Arran Reader and David Scrivener for their valuable comments on the manuscript, and to Aimee Duffus for her help with data collection. Thank you also to Maximilian Zangs for his help with the initial version of the experimental paradigm script. This research was funded by the EPSRC, EP/1503705 DTG 2014/2015 ‘Method development for coupled EEG-fMRI’.

## Conflict of Interest Statement

The authors declare that the research was conducted in the absence of any commercial or financial relationships that could be construed as a potential conflict of interest.

## Data Accessibility Statement

The raw data, pre-processed data, and analysis scripts can be found on the Open Science Framework: https://osf.io/thdva/, DOI 10.17605/OSF.IO/THDVA

